# A practical evaluation of sampling filtration and preservation of environmental DNA samples for the water column monitoring

**DOI:** 10.64898/2026.06.18.733101

**Authors:** Thierry Baussant, Adriana Krolicka, Grethe Kjeilen-Eilertsen, Thomas Merzi

## Abstract

Offshore industry still largely relies on traditional approaches for regulatory compliance to environmental impact on the water column. Implementing environmental DNA (eDNA) workflow can offer several advantages, but early stages such as sampling and conservation of the samples require standardization and simplification before they can be routinely applied in offshore monitoring programs. In this study, we assessed the effect of several filter types (Durapore disc, Sterivex capsule and Wattera high-capacity capsule; all with 0.22 µm pore size) allowing for different volume of filtration used for sampling eDNA. We also evaluated the effect of 25 days conservation of unfiltered water samples with different preservative solutions (Benzalkonium chloride -BAC, Longmire’s solution LONGI and a modified Longmire’s solution without SDS, LNoSDS) as a viable option when immediate filtration and cold storage are not possible. For downstream eDNA evaluation of filter types and preservation, we used quantitative digital PCR on selected target DNA and metabarcoding for qualitative assessment of marine prokaryotic and eukaryotic communities. Overall, filter choice had relatively less effects on quantitative and qualitative information from eDNA compared with water preservation. Sterivex and Durapore were better filter choices for biodiversity assessment. While the Wattera filter allowed processing of larger water volumes and improved quantification of metazoan DNA, handling and processing were more challenging. For water conservation, LNoSDS was the best option. Chemical agents of LONGI and BAC may provide favourable substrates for some tolerant bacterial strains, altering the microbial community composition, with consequences for the overall qualitative evaluation of conserved eDNA. For targeted metazoan eDNA, however, chemical preservation showed clear benefits. This research highlights key considerations and viable options for eDNA sampling and simple preservation workflows without cold storage for implementation in offshore water column monitoring.

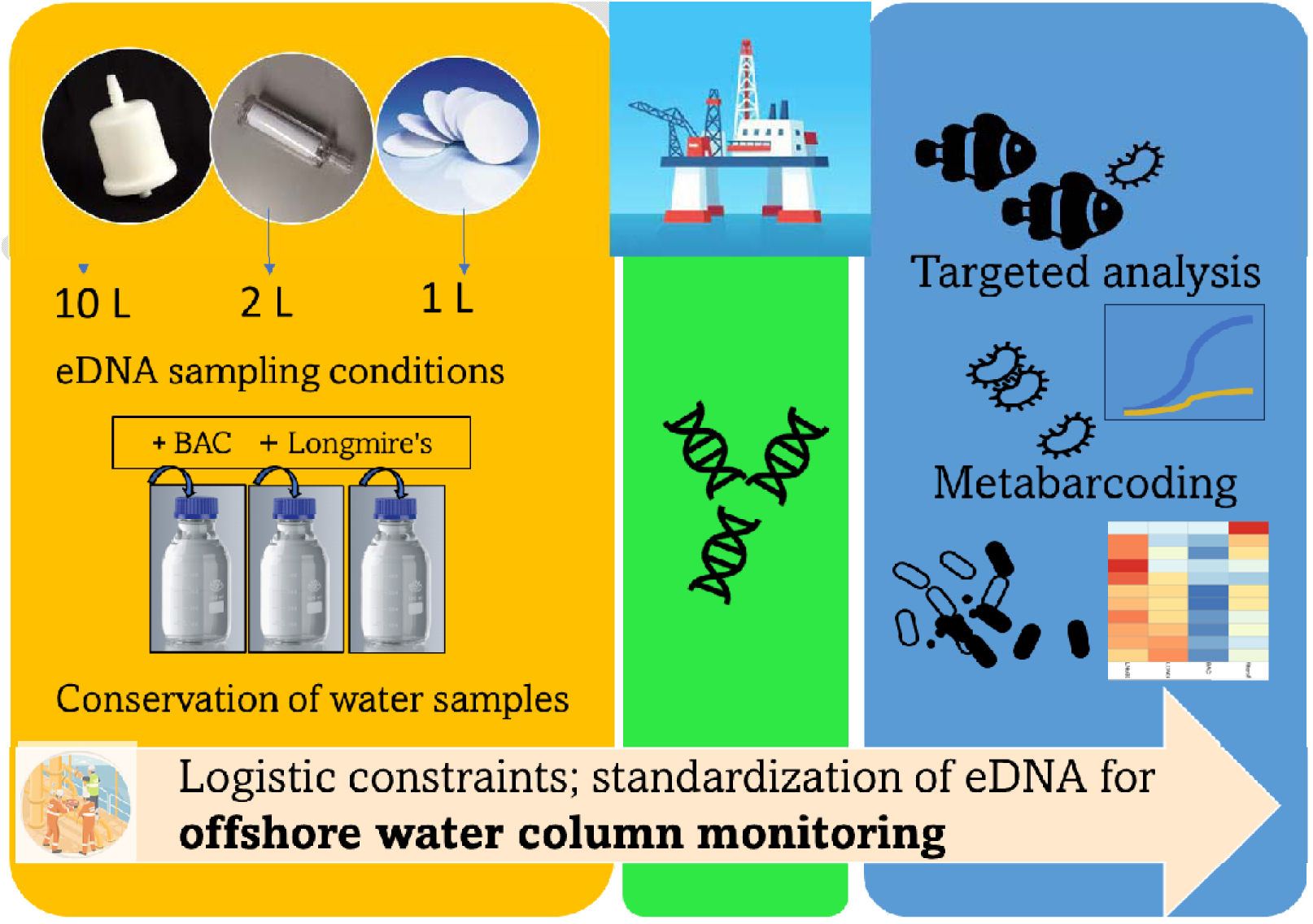

**Highlights:**

- Need for standardization of eDNA workflow for offshore water column monitoring
- Importance of eDNA sampling (filters) and eDNA conservation (preservatives)
- Filter choice does not affect drastically the dominant eDNA communities
- Conservation outside cold storage challenging for eDNA-based biodiversity evaluation
- Viable options: Sterivex filter for sampling; Longmire’s (no SDS) for conservation

## 1. Introduction

To monitor the offshore water column and evaluate the impact of offshore effluents on the marine environment, environmental DNA (eDNA) could be a valuable alternative to complement or replace the current approaches based on risk of exposure and effects (biomarkers) measured in selected field biological elements (wild fish, caged bivalves). For more than a decade, the use of eDNA has revolutionized the way species can be monitored in their environment (Deiner et al., 2017), including the application in ecological assessment for marine species diversity assessment (Andruszkiewicz et al., 2017; Djurhuus et al., 2018; Govindarajan, 2023), impacts assessment (Greco et al., 2022; Wang et al., 2019) and for management purposes (Kelly et al., 2014). eDNA allows for the detection and quantitative evaluation of the presence of specific taxonomic targets related to chemical pollution, pathogens or invasive species invasion by using sensitive nucleic assays (qPCR, dPCR, and ddPCR) at a fraction of the cost and time used by traditional methods. Furthermore, with eDNA metabarcoding, one can address exhaustive inventories about all organisms present in the marine environment, including those not conventionally used in biomonitoring such as microbial and meiofaunal taxa. This provides a broad and ecological-relevant evaluation of the biodiversity at an ecosystem level valuable for environmental managers and stakeholders involved in ecosystem-based assessment. Despite these advantages, eDNA-based methods are still not fully integrated into a regulatory framework (Darling, 2020) but more recently specific and generic guidelines for eDNA water research and workflow for different environment and regions have been published (Altermatt et al., 2023; De Brauwer et al., 2023; Gold et al., 2022; Minamoto et al., 2021; Oladi et al., 2024; Sanches and Schreier, 2020). There are also an increasing number of studies using eDNA in the context of impact assessment related to offshore industry, and more so for compliance monitoring of benthic communities around platforms (Krolicka et al., 2020; Lanzén et al., 2021; Laroche et al., 2018; Mauffrey et al., 2021; Pawlowski et al., 2024), but to a lesser extent in the water (Alexander et al., 2023; Alexander et al., 2022; Cordier et al., 2019; Hestetun et al., 2023). These aim to evaluate species assemblages and support decision-making for seafloor drilling operations, infrastructure presence or water effluent discharges management. For water column around offshore oil and gas infrastructure, recent studies demonstrated the versatility of this approach (Alexander et al., 2023), but this is still limited and today more traditional methods are used for environmental assessment (Beyer et al., 2025; Beyer et al., 2020). Companies operating on the Norwegian continental shelf are required to carry out every 3 years environmental monitoring to obtain information on the actual and potential environmental impacts of their activities and to provide authorities with a better basis for regulation (Norwegian Environment Agency, 2023) (Cordes et al., 2016). Further, the operators shall, as part of the environmental monitoring, contribute to developing new methods for monitoring the water column. It is suggested to monitor of several representative species with different ecological functions and from several trophic levels, where eDNA is expected to provide useful information for future WCM (Norwegian Environment Agency, 2023) (Bravo et al., 2023). Yet, before eDNA-based methods can be used in offshore field survey, adequate qualification and standardization of eDNA workflow need to be provided in guidelines to support routine monitoring (Hinz et al., 2022). Resources and facilities available for offshore field sampling can be very limited. The most common way to capture eDNA is often relying on sampling, filtration and then preservation of filters either dry, frozen or in lysis buffer. Sampling requires some water collection device such as Niskin container. Filtration requires a water pump and adequate filter, a key component of the sampling workflow. Then the optimal water volume and corresponding filter type are selected based on the target taxonomic group, and the specific research objectives of the monitoring. Immediate freezing of samples after filtration is an ideal approach but require access to a cold storage chain. Alternatively, simple water samples can be collected and conserved during routine operation until time and logistic allow to process them by personnel in the laboratory. Evaluating these early steps of eDNA workflow are needed to facilitate their routine integration in offshore operations (Majaneva et al., 2018; Patin and Goodwin, 2023; Takahashi et al., 2023).

### DNA filtration and volume

Despite much effort to optimize eDNA laboratory analysis, the impacts of eDNA capture methods are underexplored. The review by Bunholi and co-workers (Bunholi et al., 2023) indicates that there is a clear lack of standardization across water filtration procedures in the eDNA studies. For the eDNA studies across the six most prevalent aquatic systems (coastal ocean, river, lake, bay, open ocean, coral reef), the median volume of filtered water was 1 L, but volumes ranged from 0.002 to 150 L, with a mean (± SE) volume of 4.74 ± 13.06 L. The optimal volume of filtered water remains unresolved in the eDNA community (Bessey et al., 2021; Bunholi et al., 2023; Hunter et al., 2019; Schabacker et al., 2020). Filtration strategy is a key aspect of eDNA surveys and tradeoffs variables include target organisms, filter porosity, volume, time, and not least resource availability. Common filters used in the field are open disc filters (such as Durapore) but enclosed filters (“capsules”) are becoming routinely used for capturing eDNA (Patin and Goodwin, 2023; Spens et al., 2017; Takahashi et al., 2020). The effective filtration area of a 47mm Durapore filter (∼13 cm^2^) is equivalent to that of the Sterivex capsules (∼10 cm^2^) and Durapore can also be housed for in-line filtration. For other applications, high-capacity capsules such as the Waterra (McAdam, 2020) offer a much larger surface area of the filter (up to 600 cm²), and can be advantageous to process large volumes of water, although these filters are currently not very much used for marine waters (but see (Roblet et al., 2024). In the offshore environment, eDNA can be more patchy and require the sampling of higher volume of water over standard volume (2L or less) (Kumar et al., 2022). Large-volume processing for eDNA collection in marine communities, while effective for patchy, rare or elusive species (Govindarajan, 2023; Govindarajan et al., 2022; Pochon et al., 2024; von Ammon et al., 2020), elevates the risk of co-extracting PCR inhibitors proportionally to the volume of organic matter filtrated and require the use of a large volume of expensive reagents for DNA extraction. To the best of our knowledge, a comparison of the performance of high-capacity capsules to lower capacity filters and capsules is still missing. On-site filtration, even if preferable (Patin and Goodwin, 2023), might not be possible in some instances, such as (1) when intensive sampling is conducted over a short time period (Takahashi et al., 2023) (2) when several research, and simultaneous sampling activities on the same vessel require task prioritization (for example mentioned in Nilssen and Bakke (Nilssen and Bakke, 2011) and also (3) when there is limited personnel and infrastructure capacity such as no cold storage. Moreover, carrying out filtration in uncomfortable and chaotic conditions also increases the risk for cross-contamination.

### Conservation outside a cold storage chain

Storage of water samples at ambient temperature or low temperature for longer periods is not advisable because of microbial activity, and eDNA degradation by exogenous enzymes (Kumar et al., 2020; Takahashi et al., 2023). When cold storage chain is lacking, the use of preservatives directly added to the sampled water can minimise microbial enzymatic activity and be one viable and field-friendly alternative. Longmire’s solution is a widely used tissue preservation solution containing different agents of which sodium dodecyl sulfate (SDS) is a strong anionic detergent that acts by breaking down cell membranes to release DNA. Using this solution, Williams et al. (Williams et al., 2016) found that freshwater eDNA was best preserved at concentration rate of 1:3 (5mL Longmire’s:15mL sample water) and with eDNA positive detection comparable to their positive control (i.e., freezing) for up to 56 days and Mauvisseau et al.(Mauvisseau et al., 2021) showed that Longmire’s solution added to Sterivex filters used to collect eDNA for 3 fish species facilitated the best recovery after 8 months. Although results with this solution are encouraging, there are also uncertainties for post-analysis regarding substantial inhibition in PCR reaction following some DNA extraction method (Tomke et al., 2025) and the use of the SDS chemical (Boiso and Hedman, 2017) that raise questions. Further, its effectiveness for marine water samples and metabarcoding analysis need to be tested (Bizzozzero et al., 2024). An alternative to Longmire’s is the addition of Benzalkonium chloride (BAC), a cationic surfactant chemical that suppresses eDNA degradation by most microorganisms, reported effective for targeted analysis in fresh- and seawater samples preserved for 10 days at ambient temperature (Yamanaka et al., 2017). Further, the findings of Jo et al. (Jo et al., 2021) also indicated that preservation with BAC does not affect marine fish species richness using metabarcoding in various environmental conditions. However, the effectiveness of BAC to preserve eDNA samples for generic biodiversity evaluation in comparison to other preservation strategies remains to be tested (Sales et al., 2019). To summarize, the use of preservatives such as Longmire and BAC can be good alternative for conservation of water samples and single species detection, but further tests are needed since there are ambiguities regarding their flaws for metabarcoding analysis and marine biodiversity assessment.

The objective of this study was to evaluate the impact of (1) filter type, and consequently, the volume of filtrated water, (2) eDNA conservation strategies on metabarcoding and targeted eDNA results. For (1), we compared the performance of commonly used filter types (Durapore, Sterivex and Waterra). For (2), we evaluated the efficiency of standard and modified (SDS-free) Longmire solution and BAC in preserving directly water samples, as field-friendly alternatives to filtration and freezing conservation. Both (1) and (2) employed 16S- and 18S rRNA metabarcoding for species richness evaluation and digital PCR for quantitative (copy number) evaluation of two taxa, metazoan (Teleostei) eDNA and unicellular eDNA (Dinophyceae). Our assumptions were that no significant difference in the community composition and quantitative eDNA output resulted from filter type, volume or seawater preservation options. The results can help offshore operators to select adequate early stages of eDNA workflow and highlight options valuable for the water column monitoring.

## 2. Material and methods

### 2.1 Sampling

Sampling of water was performed in March 2021 around an offshore field (56°32.477N, 3°12.684E) in the North Sea. A high-flow BP 1301 water pump (Sweden) was used first to pump water from 10 m below surface to 25L plastic tanks onboard the ship. The water was collected at one station approximately 750m away from the platform. On board the ship, water was either filtrated immediately using different type of filters (see below 2.2) or preserved in 1L glass bottles using different buffer solutions (see below 2.3). Six replicate samples were collected for all samples except Sterivex for which only three were collected (Figure 1).

**Figure 1.**
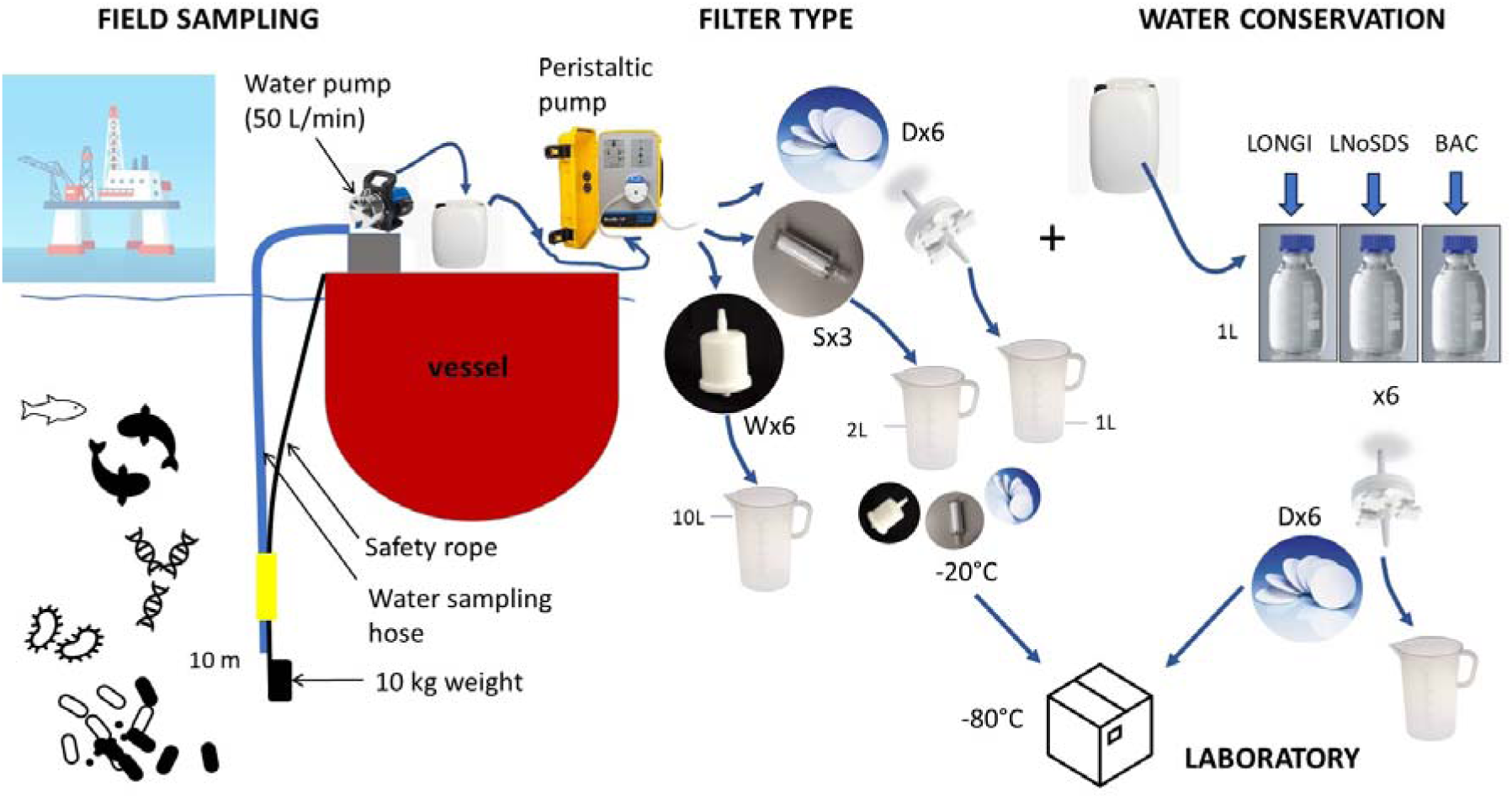
Overall flowchart of the offshore water sampling and processing of the samples in the present study.

### 2.2. Effect of filter type and water volume on eDNA recovery

The effect of several filter types and water volume on eDNA capture was investigated for the 6 replicate samples collected at the sampling station. Wattera (PES, 0.22 µm, Waterra USA Inc.) and Sterivex-GP (PES, 0.22 µm, Merck Millipore, Burlington, Massachusetts, USA) capsules were used for the collection of 10L and 2L of seawater, respectively. Also, Durapore disc filters (PVDF, 0.22 µm, diameter 47 mm, Merck Millipore, Burlington, Massachusetts, USA) were used to collect 1L water. For capsules and disc filters, water was pumped from the 25L plastic tanks using a Masterflex L/S portable Cole-Parmer (Cole-Parmer Instrument Company Ltd, London, England) sampling pump. For the capsule filters, a large 50 ml syringe was used to remove the residual water from the filters. For disc filters, a polypropylene in-line filter holder (Merck Millipore, Burlington, Massachusetts, USA) was used to fit a disc for each replicate sample. 6 replicate samples were collected for each filter type except for Sterivex where only 3 were collected. Both capsules and disc filters were stored at −20°C onboard the vessel then at −80°C after arrival to the laboratory.

### 2.3. Water conservation

On the ship, 1L Duran glass bottles were prepared with three different water conservation solutions: Longmire solution -LONGI (Longmire et al., 1997), Longmire solution without SDS -LNoSDS and Benzalkonium chloride -BAC (Yamanaka et al., 2017). For LONGI and LNoSDS, we used a 1:5 ratio of buffer to water. For BAC, the concentration was 0.01% as used by others (Kumar et al., 2020). The samples were conserved onboard 4 days at air temperature. In the laboratory, the samples were conserved another 3 weeks at room temperature before processing. Then, each replicate bottle with conserved water was filtered on a 47mm disc Durapore filter fitted to an in-line filter polypropylene holder (MilliporeSigma™,USA) with a 7.6 cm diameter and with a peristaltic pump (Watson Marlow, UK). Between each solution, the water line and the in-line filter holder were rinsed using Millipore water. Six replicate 1L glass bottles were used for each conservation method. Following filtration, the filters were stored at −80°C in the laboratory. As reference to analyse the effect of conservation on targeted species and species composition, we used the Durapore frozen samples (Durapore; table 1).

**Table 1.** Overview of the samples, sampling method, sampling conservation, type of filters and volume filtered used for eDNA evaluation in this study. ^(1)^ Filters conserved at-20°C onboard the vessel during the survey (4 days) and −80°C in the laboratory; ^(2)^ Water samples conserved onboard the ship for 4 days at air temperature then a further 3 weeks in the laboratory at room temperature before filtration and storage at −80°C.

| N replicates and sample name | Sampling method | Sampling conservation | Filter type | Volume of water |
| --- | --- | --- | --- | --- |
| **N=3; Sterivex** | water collection and immediate filtration | Freezing at -20 then -80°C <sup>(1)</sup> | Sterivex/ polyethersulfone (PES) 0.2 µm/10cm<sup>2</sup> | 2L |
| **N=6; Durapore** | water collection and immediate filtration | Freezing at -20 then -80°C <sup>(1)</sup> | Durapore/Polyvinylidene fluoride (PVDF) 0.2 µm/17cm<sup>2</sup> | 1L |
| **N=6; Waterra** | water collection and immediate filtration | Freezing at -20 then -80°C <sup>(1)</sup> | Wattera/ polyethersulfone (PES) 0.2 µm/600cm<sup>2</sup> | 10L |
| **N=6; LONGI** | Water collection only | Addition of Longmire solution to water <sup>(2)</sup> | Durapore/Polyvinylidene fluoride (PVDF) 0.2 µm/17cm<sup>2</sup> | 0.8 -1L |
| **N=6; LNoSDS** | Water collection only | Addition of Longmire solution without SDS to water <sup>(2)</sup> | Durapore/Polyvinylidene fluoride (PVDF) 0.2 µm/ 17 cm<sup>2</sup> | 0.8 -1L |
| **N=6; BAC** | Water collection only | Addition of BAC solution to water <sup>(2)</sup> | Durapore/Polyvinylidene fluoride (PVDF) 0.2 µm/17 cm<sup>2</sup> | 0.8 -1L |

### 2.4. DNA extraction

Sterivex and Durapore filters: Sterivex filters were opened by cutting the end using a sterilized PVC tube cutter, the filter was removed from the casing using sterilized forceps. This step was performed on a sterile Petri dish (Cruaud et al., 2017). For disk filters (Durapore), this step above was omitted. Afterwards the filter pieces (from Sterivex capsule and Durapore) were transferred into 2 mL Eppendorf tube. The 3M guanidine thiocyanate (1,5 ml) lysis solution pH 8.9 (3M GuSCN, 50mM Tris, 15mM EDTA. 2% Sarkosyl and 0.2% SDS) was added and the samples were incubated at −85 C for 10 min, the incubation was performed with a slight rotation. After filtration of lysate through a Millex-GV syringe filter (Millipore, pore size: 0.2□μm; diameter: 13□mm) 1000□μl of sterilized lysate and 900□μl of diluent (555□mM sodium acetate pH□5.2 in 70% ethanol [v/v]) were combined. Afterwards, the entire volume of the lysate was passed through a spin column in the DNeasy Tissue Kit (Qiagen, Hilden, Germany) and washed twice before DNA was eluted with an 80□μL elution buffer. After extraction, the concentration of DNA in each sample was measured using the Qubit dsDNA HS (High Sensitivity) Kit (Thermo Fisher Scientific, Carlsbad, California, USA) before the samples were stored at −20 °C.

#### Wattera filters

When the water filtration was completed 50 ml syringe was used to remove remaining water in the Wattera capsule. 35 ml 3M guanidine thiocyanate lysis solution pH 8.9 (3M GuSCN, 50mM Tris, 15mM EDTA. 2% Sarkosyl and 0.2% SDS) was added to capsules (from the narrow part) using 10 ml automatic pipette. The filters from both sides were closed (but not too tight) with the caps provided by the supplier. The filters were placed on the glass beakers and incubated at 85 °C 20 min, wherein samples were vortexed 3 times in between. The lysate was transferred to 2 falcons. One volume of phenol:chloroform:isoamyl alcohol (25:24:1) was added to the sample, and vortexed for approximately 20 seconds. The samples were centrifugated at RT for 40 minutes at max speed (5000 rpm). The upper layer was transferred to falcons and 1 volume of solution Isopropanol -sodium acetate to the final concentration of 0.3 M was added. After overnight incubation at RT., the samples were centrifugated at RT for 60 minutes at max speed. The supernatant was discarded but 2-3 ml of supernatant was kept. The extract was loaded on the commercial spin columns provided in DNA Blood and Tissue kit (QIAGEN). Finally, 2 steps of ethanol washing were performed as described in the DNA Blood and Tissue kit manual. DNA was recovered from the column using 100 µl of elution buffer provided in the QIAGEN kit. The concentration of recovered DNA in each sample was measured using the Qubit dsDNA HS (High Sensitivity) Kit (Thermo Fisher Scientific, Carlsbad, California, USA) before the samples were stored at −20 °C.

### 2.5. Targeted analysis using digital PCR (dPCR)

Targeted amplification of 18S rRNA gene copy number of Dinophyceae (Krolicka et al., 2020) and 12S rRNA mitochondrial gene copy of Teleostei (Taberlet et al., 2018) was investigated to quantify marker gene copies (MGC) from the different filters or following the different preservation solutions. This was performed on an QIAcuity One instrument (QIAGEN; Hilden, Germany) using QIAcuity Probe PCR Kit or QIAcuity EvaGreen (EG) PCR Kit respectively according to the manufacturer’s instructions. Oligonucleotides were added to a final concentration of 500 nM for primers and 250 nM for probe (Supplementary Table 1.). Samples were manually loaded into the QIAcuity™ 26K 24-well Nanoplate (QIAGEN, Hilden, Germany). Amplification steps were carried out in accordance with the manufacturer’s thermal protocol. Partitioning, processing, and fluorescent signal processing was automatically carried out by proprietary software (QIAcuity® Software Suite Version 1.2).

### 2.6. Metabarcoding: amplicon generation and library preparation

Metabarcoding analysis was performed with the services of Novogene (UK) Company Limited. PCR amplification of targeted regions was performed by using specific primers connecting with barcodes. Amplicons for 16S rRNA genes were generated using the primers pairs 341F and 806R (Yu et al., 2005) targeting the hypervariable regions V3-V4. For 18S rRNA, the primers pairs targeting the hypervariable V4 region were 528F and 706R (Cheung et al., 2010) (Supplementary Table 1). The PCR products with proper size were selected using 2% agarose gel electrophoresis. The same amount of PCR products from each sample was pooled, end-repaired, A-tailed and further ligated with Illumina adapters. Libraries were sequenced on a paired-end Illumina platform to generate 250bp paired-end raw reads. The upgraded NovaSeq sequencing platform was implemented for the PE250 strategy of paired-end sequencing.

### 2.7. Bioinformatics

Paired-end reads were assigned to samples based on their unique barcodes and truncated by cutting off the barcode and primer sequences. The QIIME 1 pipeline was utilized for the analysis of rRNA gene sequencing data. Initially, raw sequencing reads were processed to obtain “Clean Tags” (also known as quality-filtered reads). Chimeric sequences within these Clean Tags were subsequently detected and removed using established algorithms, resulting in “Effective Tags”. Operational Taxonomic Units (OTUs) were then generated by clustering these Effective Tags at a 97% sequence identity threshold. Taxonomic annotation of the representative sequences for each OTU was performed against the Silva 138.2 ribosomal RNA gene database. According to OTUs annotation results and sample characteristic tables, species abundance tables at different taxonomy levels were generated and used for the further statistical analysis.

### 2.8. Statistics and data visualization

For dPCR analysis, GraphPad Prism version 5.0 was used (GraphPad Software, San Diego, California USA) to test for normal distribution of the data and further analyse them for significant differences using either ANOVA or a two-tailed Mann Whitney test. Significance was set to a level of 0.05. Regarding metabarcoding, OTUs abundances were first normalized and used for the subsequent analysis of alpha- and beta-diversity. First, the top 10 taxa of each sample at the genus level were selected to plot distribution histograms (Barplots) of their relative abundance in Perl (version 5.26.2) through SVG function. Venn diagrams were produced in R (Version 4.0.3) using the VennDiagram() function to analyze both the common and unique OTUs for the different samples, and the Chao1 richness index was calculated for alpha diversity analysis with QIIME (Version 1.9.1) and displayed with R. Differences in Chao1 between groups were analyzed using a Kruskal-Wallis test. Further, beta diversity analysis was performed to evaluate the differences between samples in OTUs assemblage. Non-metric multidimensional scaling (NMDS) was used for data dimension reduction and was implemented through R software with ade4 package and ggplot2 package. Further, the abundance information of the top 35 taxa of each group at the genus taxonomic rank was used to draw heatmap, visually displaying similarity and differences in samples, using the pheatmap() function in R. The unifrac distance between samples was realized in Perl for checking the samples clustering. For analysis of the community differences between samples, LEfSe (linear discriminant analysis (LDA) Effect Size) was performed to detect characteristic taxa and potential bioindicators. LDA scores larger than the set threshold (4 set by default) were considered as significantly different.

## 3. Results and Discussion

### 3.1. Evaluation of the effect of filter type and the sampling volume on eDNA analysis

#### DNA yield and targeted quantities

Before volume normalisation, total DNA yield was significantly higher in the water samples collected with Wattera. However, after normalisation to 1l, there was no significant difference in DNA yield between the different filters (Supplementary Figure 1A). Further, before volume normalization, MGC (marker gene copy) of Teleostei and Dinophyceae was significantly higher for the Wattera filters (Figure 2). After normalization to volume, MGC of Dinophyceae was not statistically different between the filters (Mann-Whitney, p>0.05) but MGC Teleostei (Fish) from Wattera filters still remained significantly higher than from Sterivex filters (Mann-Whitney, p<0.001). In addition to filter type, which determines water flow rate and effective filtration area, concentration, distribution and states of eDNA for different species can affect the quantity of eDNA recovered from the water. Compared to unicellular eDNA (such as for Dinophyceae), metazoan eDNA (such as fish) exists in a range of states (e.g., dissolved, particle adsorbed, intracellular, and organellar), and the relative distribution of eDNA among the different states requires larger volume to increase the probability of detection (Mauvisseau et al., 2022). Furthermore, eDNA from Durapore and Sterivex filters were extracted using the same DNA extraction protocol, but that differed for Wattera filters for which a phenol-chloroform-isoamyl (PCI) method was used. Substantial inhibition was observed in preserved samples when using a PCI extraction method (Kumar et al., 2022; Tomke et al., 2025). In addition to volume of water, type of filters and the range of eDNA state distribution for the different species, there is a possibility that the PCI step used to extract eDNA from Waterra filters acted differently on the MGC of the two taxa. This underlines the complexity of multi-trophic studies and the need for best practices and standardization regarding the analytical workflow (i.e., filter, capture, preservation, extraction, and detection methods) (Goldberg et al., 2016; Patin and Goodwin, 2023).

**Figure 2.**
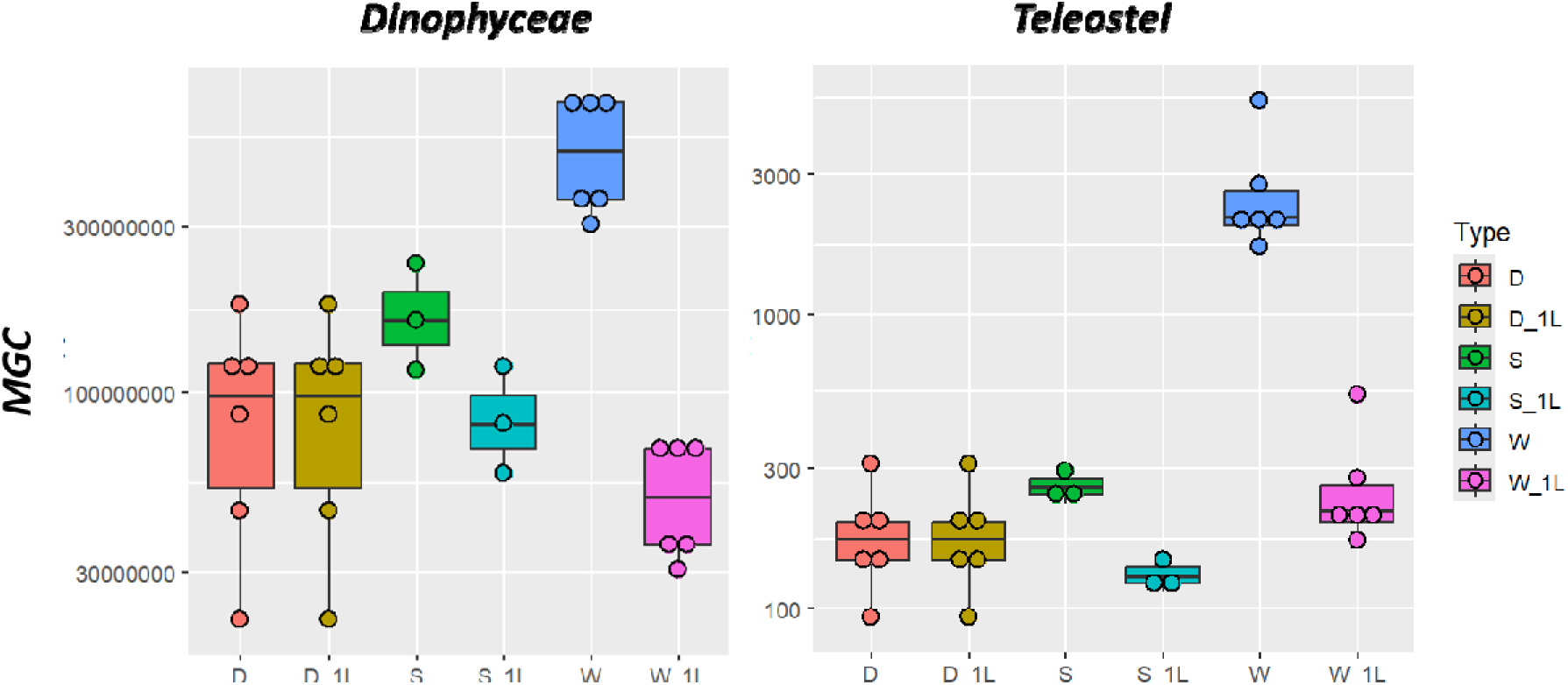
Targeted eDNA quantification with filter types.

#### Metabarcoding

##### Prokaryotes

A total of 2633 OTUs were identified in the dataset and 1203, 1927, and 1310 OTUs were associated to Sterivex, Durapore and Waterra, respectively. The relative abundance of the 10 dominant taxa was quite similar for all types of filters with a major contribution from cyanobacteria accounting for 40 to 50% of all taxa (Figure 3A). Noteworthy is also the relatively high abundance of archea taxa from the Marine_Group_II clade, and Nitrosopumilaceae (with *Candidatus_Nitrosopumilus* as a dominant species) which have been associated to hydrocarbons degradation (Knapik et al., 2020; Peeb et al., 2022; Rinke et al., 2019; Zhang et al., 2015). Of the OTUs detected from these filters, a relatively high number (approximately half) were common to all, and Durapore showed the highest number of unique OTUs (Venn diagram, Figure 3B). Despite a higher total DNA yield from Waterra samples (Supplementary Figure 1A), 16S-OTUs richness characterized by the Chao1 index was significantly lower in these samples compared to Sterivex and Durapore (Figure 3C). Further, the beta-diversity NMDS plot showed some spreading within Waterra and Durapore but overall prokaryotic OTUs assemblage from the different filters appeared relatively close Figure 3D). The observed spreading on the NMDS plot may be explained in differences revealed by the heatmap plot (Supplementary Figure 2A), which visually displayed different abundance and clustering for several taxa, and the LefSe (Supplementary Figure 3A), which revealed significant differences in the relative abundance of OTUs and identified a few specific taxa characterizing the different samples. Whilst marine group II euryarchaea characterized Waterra samples, OTUs from Clade_Ia, Clade_II and SAR86_clade (Alphaprotebacteria) were rather identified as characteristic features for Sterivex and Durapore. Furthermore, genus like *Colwellia* and *Pseudoalteromonas* (Gammaproteobacteria), which are typical oil-degrading bacteria, were relatively more abundant in Durapore. From the location of this study around an offshore field, it is not surprising to detect these genera, but it is noteworthy. Studies have shown that 0.22 μm Durapore filters are well suited to the recovery of hydrocarbonoclastic bacteria (Knapik et al., 2020; Krolicka et al., 2017; Krolicka et al., 2019).

**Figure 3.**
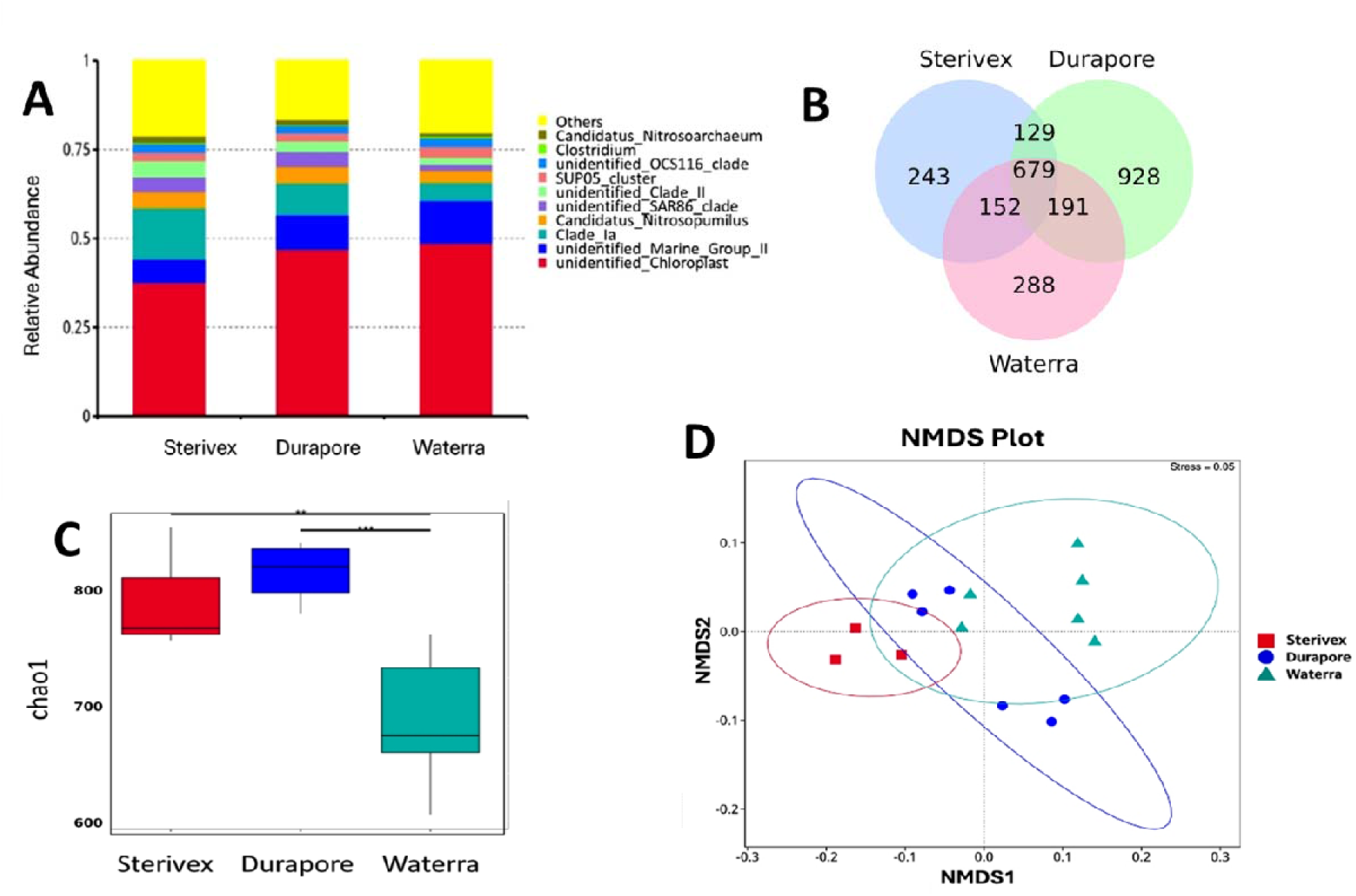
eDNA metabarcoding of Prokaryotes recovered from different filters-

##### Eukaryotes

As much as 2345 OTUs were identified totally from the different filters used to collect eDNA and a relatively equal number of OTUs were associated to Sterivex (1036), Durapore (1366) and Waterra (1375). The two main dominant taxa were represented by Calanoida and Mediophycea (diatoms) each representing from ∼10 to 25% of the total OTUs abundance. Several Chloroplastida taxa were also well represented in all samples but mostly in Sterivex and Waterra (Figure 4A). Of the total OTUs identified, there were a relatively large fraction (i.e. half or so) that were common to all samples but Durapore and Waterra showed the highest number of unique OTUs compared to Sterivex (Venn diagram, Figure 4B) and there was no significant difference in the chao1 index (Figure 4C). The 18S-OTUs community profile was relatively consistent and OTUs identified from the different filters overlapped to a relatively large proportion but with some spreading for Waterra (NMDS plot, Figure 4D). Heatmap and LeFSe identified however differences in community assemblage such as Calanoida and Chloroplastida significantly more prominent in Sterivex, Mediophycea (diatoms) but as well Trachymedusae in Durapore and, in Waterra, significant prominence of Spirotrichea from the Ciliophora phylum and occurrence of a relatively larger number of OTUs belonging to Appendicularian Oikopleuridea from the Tunicata phylum (Supplementary Figures 2B and 3B).

**Figure 4.**
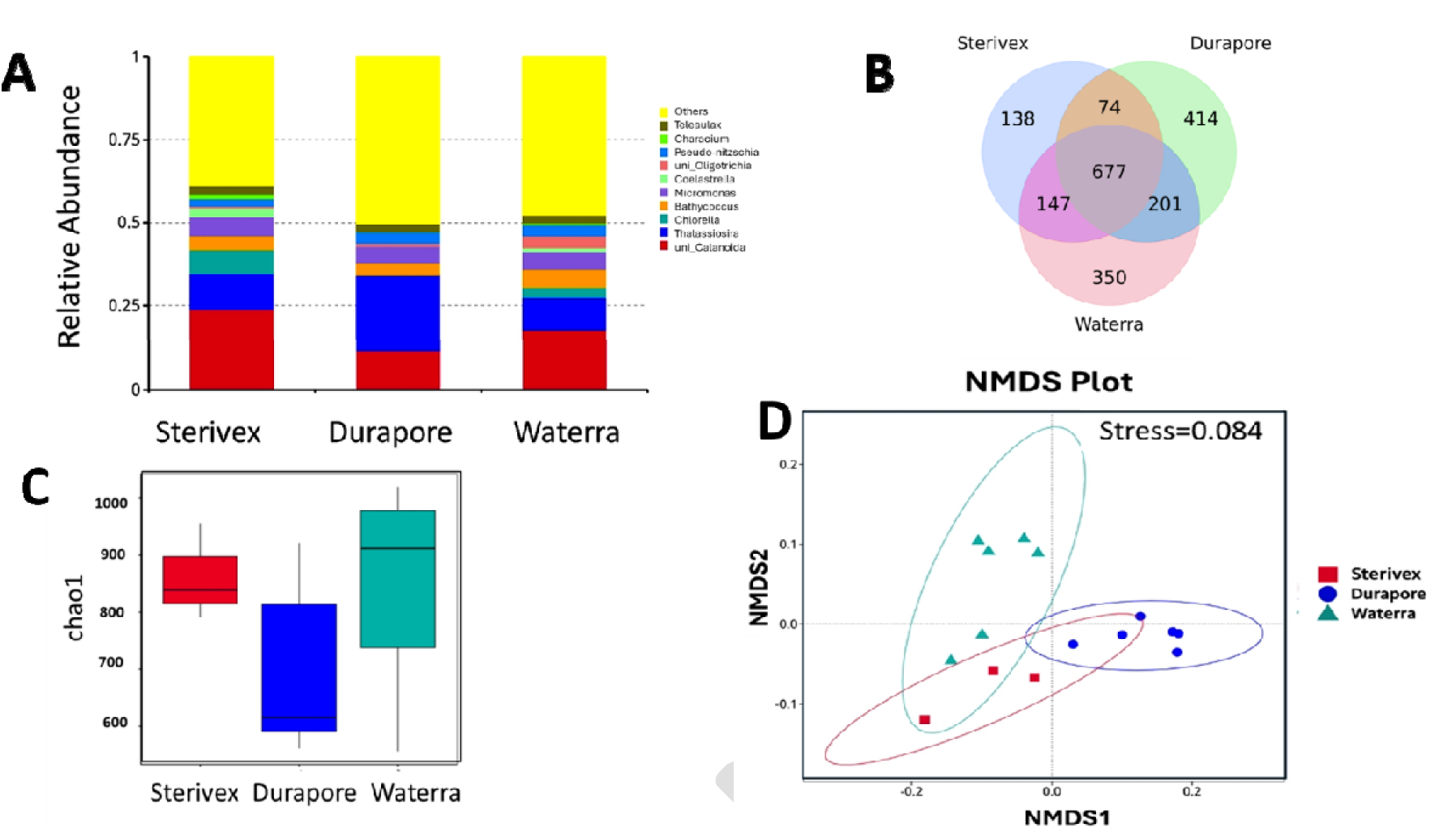
eDNA Metabarcoding of Eukaryotes recovered from different filters

Both filter type and sample volume are important parameters that can influence target taxa detection and community composition metrics. Although some differences in species assemblages were observed by using the different preservative methods, there was a high fraction of common 16S-rRNA and 18S-rRNA OTUs in all treatments. Increasing sample volume could result in increase species richness in Waterra but this was not observed despite a higher total DNA yield with this preservative (Supplementary Figure 1A). Species richness in Waterra was lower for prokaryotes but equal for eukaryotes than the other filters. This observation could be attributed to a better recovery of eukaryotic eDNA related to different physical state and distribution of the 18S rRNA nuclear marker encompassing a broad range of phyla (from unicellular to metazoan). We emphasize also the challenge associated with Waterra filters that needed a large volume of reagents to cover the filter surface area, which lead us to choose the PCI method for DNA extraction, less expensive, but deviating from more “standard” kit and manufacturer’s procedure used for Sterivex and Durapore filters. It is possible that the PCI method was affecting DNA integrity during extraction and suffered from PCR inhibition that resulted in the loss of low abundant OTUs in the final metabarcoding output of the Waterra samples. Overall, the benefits of increasing the volume of filtered water for broad taxa detection is more critical for low abundant or elusive taxa, or when organism’s eDNA exists in a range of states, is distributed unevenly in the water and bounded to particular matter (Mauvisseau et al., 2022; Pochon et al., 2024; von Ammon et al., 2023) or for some oligotrophic ocean environments (Govindarajan et al., 2021; Govindarajan et al., 2022; Kumar et al., 2022).

### 3.2. Evaluation of the effect of water conservation on eDNA analysis

#### DNA yield and targeted quantities

Overall, seawater samples collected on Durapore filters and frozen on the ship (Durapore, table 1, referred as FiltersFrozen in the following) and the preserved samples with LNoSDS had higher yield, but absolute DNA yield was highest in FiltersFrozen, and the lowest amount was obtained for BAC and LONGI (Supplementary Figure 1B). DNA yield obtained from the different replicates used for LNoSDS was more variable but overall gave significantly higher DNA amount compared to LONGI. Frozen and Longmire’s preserved samples have been shown to give higher yield compared to other preservation methods (Tomke et al., 2025) but variability in DNA yield in samples conserved with Longmire’s was also reported by Williams et al. (Williams et al., 2016), who attributed this to the inherent heterogeneity of DNA distribution in the water collected. However, another explanation may be the slightly more challenging process at the filtration stage with samples conserved with Longmire’s solution compared to BAC, and the influence of some agents of that solution on the DNA lysis and total DNA recovery, especially for the standard Longmire solution (with SDS). Further, for *Dinophyceae*, MGC in LONGI and LNoSDS preservations were not different to FiltersFrozen, but MGC in BAC was significantly lower (Mann-Whitney, p<0.05). Also, compared to FiltersFrozen, MGC numbers of *Teleostei* (fish) eDNA was higher following the different water preservation methods but only significantly different in LONGI and LNoSDS (Mann-Whitney, p<0.05) (Figure 5). Several studies using preservative agents indicate that in some instances preserved samples can favor the release of more DNA from the lysed cells compared to frozen samples (Majaneva et al., 2018; Williams et al., 2016). However, this process may also depend on the physical eDNA states (i.e. dissolved, particle adsorbed, intracellular, and organellar), which also differ for different taxa (Mauvisseau et al., 2022). Given the results with the Longmire buffer for targeted species quantification, this storage option could be part of a protocol for swift monitoring of a few target species using simple portable PCR device (Marx, 2015), once a set of bioindicators are identified.

**Figure 5.**
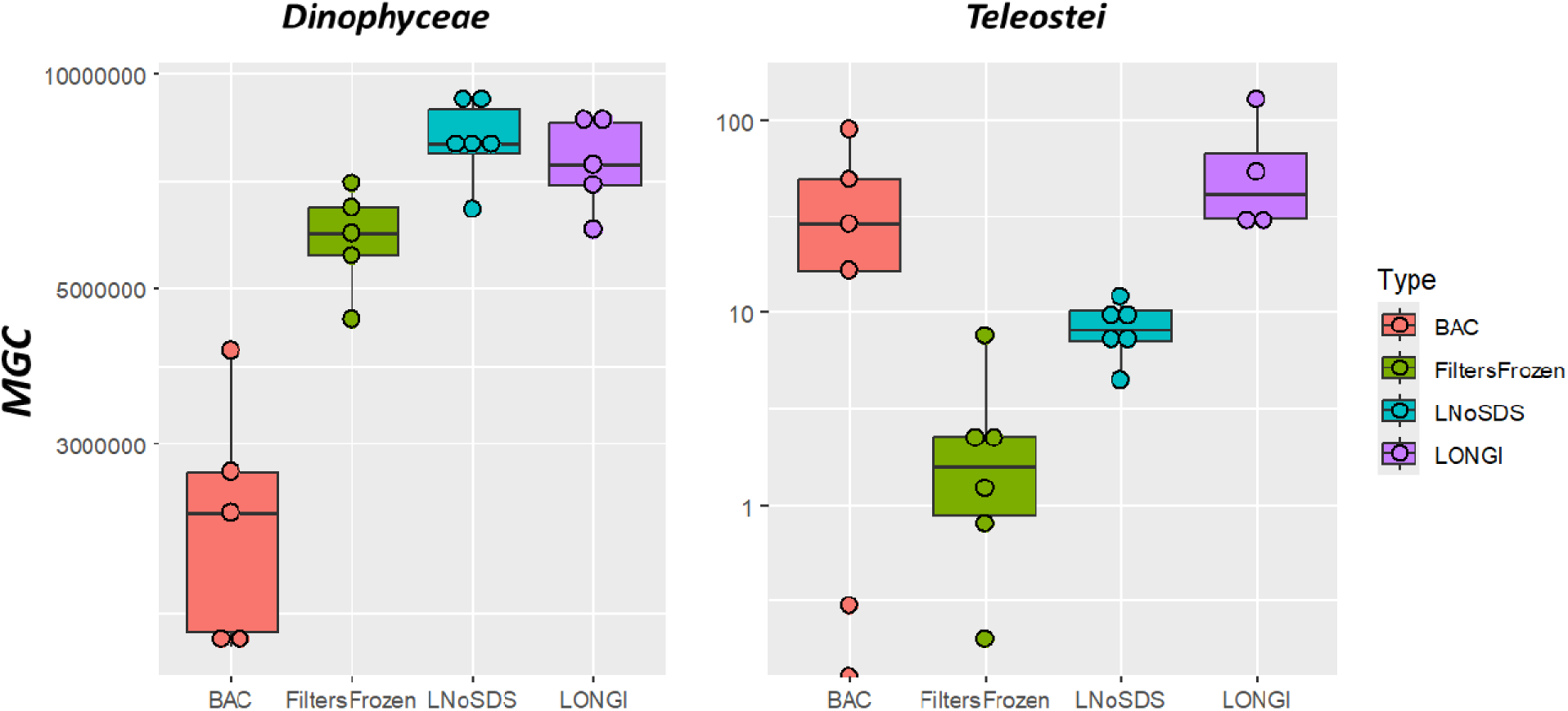
Targeted eDNA quantification for different conservation solutions.

#### Metabarcoding

##### Prokaryotes

A total of 2897 OTUs were identified in the dataset and 996, 1459, 263 and 1549 OTUs were associated to the samples preserved with LNoSDS, LONGI, BAC and filtersFrozen, respectively. The relative abundance of the 10 dominant taxa was similar in LONGI, LNoSDS and filtersFrozen (Figure 6 A) with cyanobacteria accounting for ∼50% of all taxa, and other major groups (Clade_I of SAR11; Nitrosopumilaceae) also well represented in these samples. BAC was very much different and with a dominant representation of Pseudomonadaceae and Marinobacteraceae both belonging to Pseudomonadales. Only 119 OTUs were common to all samples (Venn diagram, Figure 6B). A relatively large number of unique OTUs (885, of which 99% with a relative abundance <=0.05%) and a significantly higher number of low abundant OTUs characterized filtersFrozen (Chao1 index, Figure 6 C) compared to LNoSDS and LONGI, whilst richness and OTUs in BAC samples were as expected lowest owing to low DNA yield from the microbial lysis action of this solution (Supplementary Figure 1B). From the NMDS plot, the beta diversity showed a clustering of LNoSDS, LONGI and filtersFrozen sample replicates (Figure 6 D), indicating that the dominant prokaryote OTUs in Longmire-based solutions and in filtersFrozen were relatively similar. Nevertheless, they were clear differences in taxa abundance (supplementary Figure 4A) and significant occurrence of bioindicators at different taxonomic level identified with the LefSe analysis (supplementary Figure 5A) between the preserved and frozen filter samples. Genus of the type *Candidatus* (Archea), *Pseudoalteromonas* (Proteobacteria) and other cyanobacteria-affiliated ones were characterizing filtersFrozen, LONGI and LNoSDS, respectively. Interestingly, only LONGI recovered a larger number of hydrocarbon degraders *Oleispira*, *Alcanivorax* and *Colwellia* (Krolicka et al., 2019; Krolicka et al., 2020). Sodium Dodecyl Sulfate (SDS), a surfactant component of the original Longmire solution is readily biodegradable by many bacterial strains (Yeldho et al., 2011), with several strains isolated from marine waters (Arora et al., 2024). The growth of genus like *Alcanivorax* and *Oleispira* may be stimulated in presence of SDS, as these genera possess enzymes such as sulfatases and monooxygenases that are capable of cleaving sulfate ester bonds and oxidizing alkyl chain. Hence, the Longmire solution with SDS could alter the composition of prokaryotic organisms, favouring some to grow. Consequently, one may argue that Longmire solution without SDS is a better preservative solution for the evaluation of prokaryotic communities. While the initial premise of using BACs for eDNA preservation for prokaryotes might seem counterintuitive, our observations indicate that some bacterial species, specifically *Pseudomonas*, *Marinobacter*, and *Moritella*, exhibited robust survival in the presence of BAC (supplementary Figure 4A). A *Pseudomonas* strain was previously documented to be highly tolerant and capable of degrading BAC concentration up to 500 mg.L^−1^ in 46h (Fortunato et al., 2018). From the perspective of eDNA preservation, this highlights a critical limitation: the time-dependent degradation of BACs by these bacterial species significantly reduces the effective duration of eDNA preservation. It is possible that within the conservation time (25 days), BAC was significantly degraded by *Pseudomonas*, and potentially also by other strains such as *Marinobacter* and *Moritella*, rendering it much less effective for long-term eDNA preservation.

**Figure 6.**
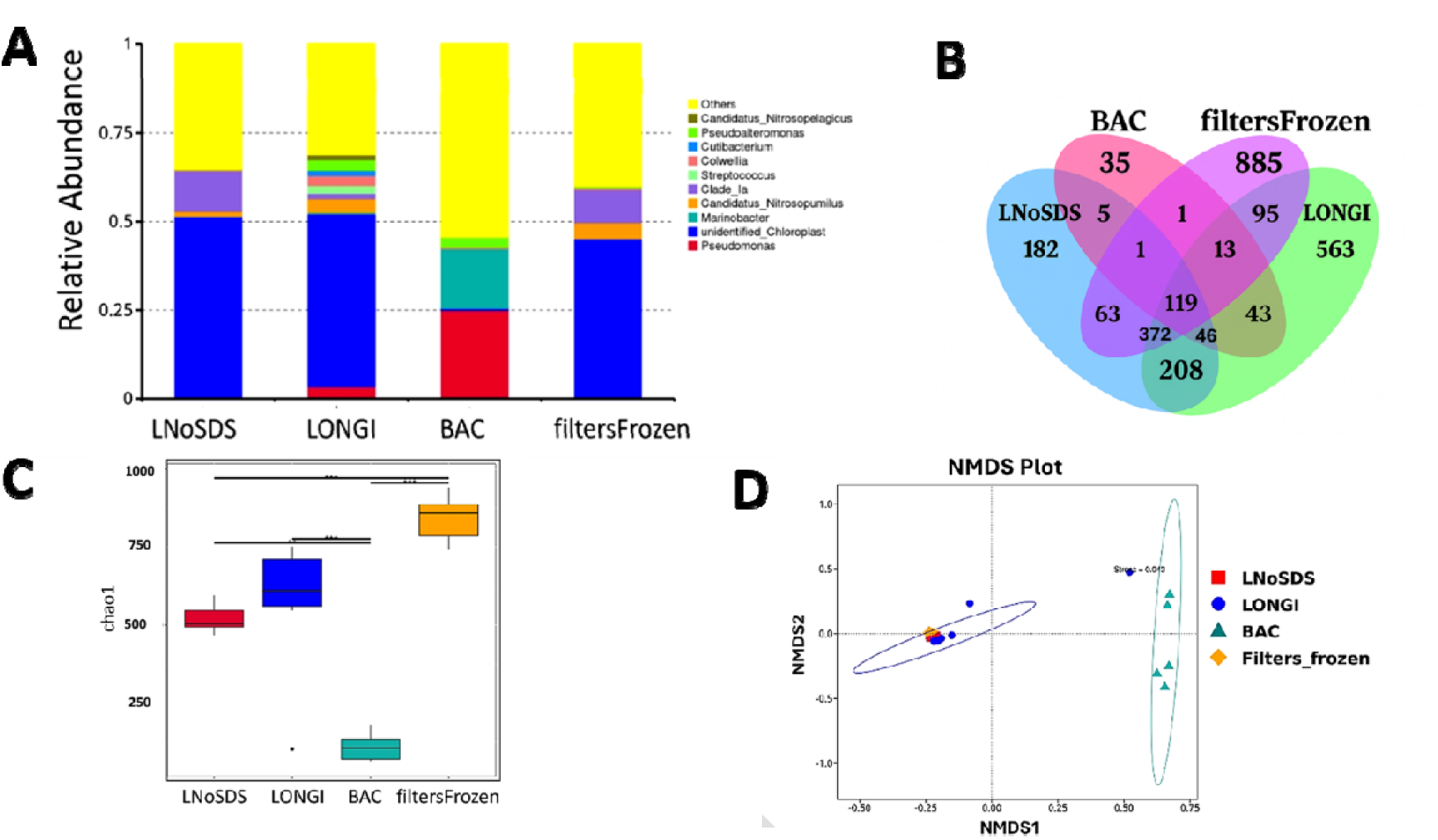
eDNA Metabarcoding of Prokaryotes recovered after 25 days conservation with different preservatives-

##### Eukaryotes

Regarding eukaryotes, a total of 2265 OTUs were identified in the dataset and approximately the same number of OTUs were present in the preserved samples (617, 987 and 730 in LNoSDS, LONGI and BAC, respectively) whilst OTUs in filtersFrozen were higher (1303). Dominant taxa such as the diatomea Thalassiosira were well represented in all samples, however relatively lower in filtersFrozen (24%) compared to BAC (34%) and LONGI (44%), reflecting the pattern observed for targeted analysis. The relative abundance of other dominant taxa (Oikopleuridae, unidentified_Malacostraca, unidentified_Calanoida) was somewhat more variable between the samples (Figure 7A) possibly related to inherent heterogeneity in eDNA distribution from larger organism, and the type of organism as depicted for targeted analysis. Generally, both the number of unique OTUs and the richness (Chao1 index) were higher in filtersFrozen compared to LONGI, LNoSDS and BAC (Figure 7B-C), also aligning with the higher DNA yield obtained from the filtersFrozen samples (Supplementary Figure 1B). The beta diversity revealed a closer similarity for water samples conserved with LONGI, LNoSDS solutions and for filtersFrozen, which were more distant to the BAC samples (Figure 7D). As for the prokaryotic organisms, there were also differences in the taxa abundance recovered from the different storage solutions (supplementary Figure 4A). The LefSe analysis identified several significantly-different characteristic taxa such as Calanoida and Chloroplastida in filtersFrozen samples, Oikopleuridea and Dinoflagellata in LNoSDS, and Malacostraca in BAC (supplementary Figure 5A).

**Figure 7.**
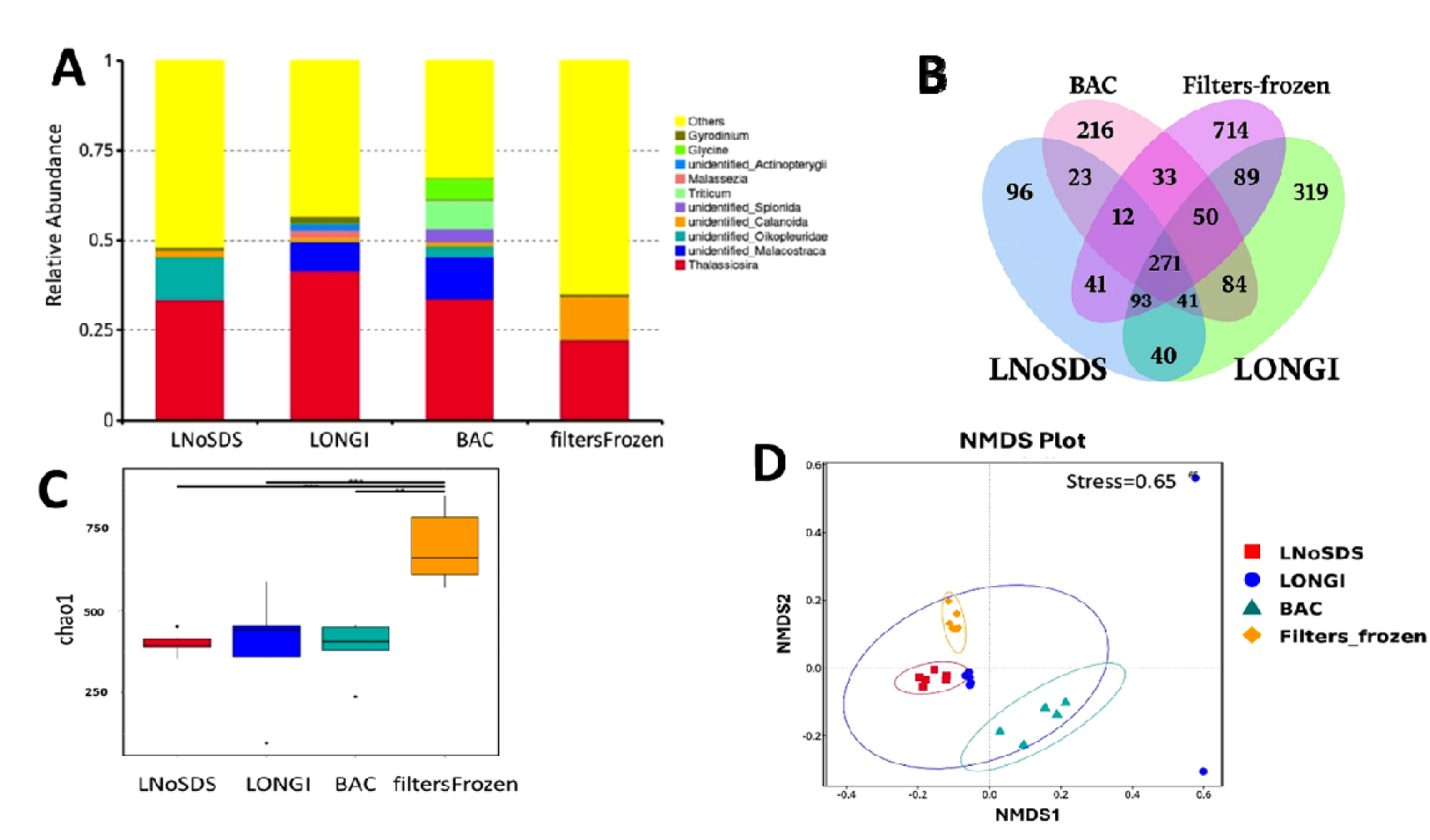
eDNA Metabarcoding of Eukaryotes recovered after 25 days conservation with different prese39

Here, we tested the use of preservatives for conservation of marine offshore samples for prokaryotes and eukaryotes community analysis using metabarcoding, which to our knowledge is new. Overall, our findings are that water conservation using preservatives is less versatile for species richness than for quantitative target species detection in the offshore environmental context of this study. However, the outcome of targeted analysis is very much dependent on the type of species and should be evaluated beforehand to select the proper conservation approach. BAC was less appropriate than the Longmire solution in preserving the quantitative signal from nuclear 18S rRNA of Dinophyceae but better for mitochondrial 12S rRNA of Teleostei. Previous studies indicate that BAC treatment substantially improve the yield and quantification of fish eDNA (Jo et al., 2021; Takahara et al., 2020; Yamanaka et al., 2017) but this is not very much studied for other species. Regarding Longmire’s. when SDS was removed (LNoSDS), total DNA yield increased but the effect on MGC for targeted analysis and on community analysis was not different compared to standard Longmire’s (LONGI). Since SDS can be degraded by several marine bacterial strains (supplementary Figures 4A and 5A), it is possible that the concentration and hence the effectivity of this agent was significantly reduced as well in LONGI, and that the two conservation treatments did not act differently in the preservation of eukaryotic eDNA. Regarding metabarcoding, the alpha-diversity of marine prokaryotes and eukaryotes OTUs in the frozen samples was significantly higher than any of the other preservation methods tested as was the total DNA yield. DNA yield is affected after 25 days conservation and that impairs the probability of detection of the low abundant species compared to freezing. Hence, our results support the general recommendation favoring the freezing of samples with minimal delay rather than using storage solutions for alpha-diversity (Patin and Goodwin, 2023; Tomke et al., 2025). Nevertheless, when LONGI and LNoSDS preservatives are used, the overall profile of the dominant prokaryotic and eukaryotic communities doesn’t seem to be modified whilst these communities in BAC are more deviating possibly owing to a reduced efficiency of this preservative over long-term. Longmire’s solution applied to filter conservation has shown to give the smallest decrease in eDNA yield and consistent results on community composition (Bizzozzero et al., 2024; Kumar et al., 2020; Mauvisseau et al., 2021). Here, this was applied directly to samples of field-collected offshore seawater, which has not been tested before. A Longmire concentration ratio of 1:5 was used to reduce chemical use and alleviate downstream analytical process in this study. Williams et al. (Williams et al., 2016) used Longmire concentrations ratio of 1:3, 1:6 and 1:15 to conserve freshwater samples and all volumes of the Longmire’s solution preserved eDNA in the samples significantly better than the “no treatment” group after 28 and 56 days. However, only the 1:3 concentration ratio was comparable to frozen sample conservation used as reference whilst the 1:6 concentration ratio preserved only half of the DNA compared to reference after 28 days. In situation where samples need to remain at room temperature for an extended period, Longmire’s buffer is an option but a concentration ratio of 1:3 seems advisable compared to 1:5 used here. Other methods of conservation have also been suggested for long-term storage such as self-preserving sampling unit or the addition of silica gel (Bowers et al., 2021; Thomas et al., 2019). Other critical considerations such as temperature fluctuations in preserved samples offshore and during transit time from offshore installation to laboratory prior to analysis need to be evaluated as well. Preservation of samples outside a cold storage chain prior to analysis is an area to investigate further for offshore managers to ease the logistic, collection and appropriate conservation of marine eDNA for monitoring.

### 3.3. Sequencing data

The sequencing data were deposited at GeneBank NCBI under PRJNA883831 BioProject

## 4. Conclusions

### 4.1. Importance of eDNA and need for protocol harmonization

Whilst environmental DNA has for a long time been used for research purposes only, other applications related to impacts from industrial use of the ocean such as for aquaculture, oil & gas production, and wind farms are increasingly being reported (Alexander et al., 2023; Alexander et al., 2022; Dahlgren et al., 2023; Hestetun et al., 2023; Peters et al., 2018; Ramey et al., 2023). The versality, evaluation for a wider spectrum of species, as well the potential for cost reduction with eDNA likely explain this. A bulk of applications is related to benthic communities and the application of eDNA for biodiversity evaluation related to compliance monitoring and regulation (Lanzén et al., 2021). In the water, a larger number of eDNA-based investigations concerns the detection of alien or invasive species (Flitcroft et al., 2025). The use of eDNA for the evaluation of targeted or general communities in the water column is also coming into more frequent use as well and this demand will likely increase in the future due to a larger use of the ocean for the blue economy (Jouffray et al., 2020) and a green energy transition where several offshore installations will also require more stringent evaluation of the impact on pelagic communities around decommissioning (Alexander et al., 2023). Hence, it is timely to implement the eDNA workflow for the offshore water column and evaluate viable options for early steps such as presented here for optimal DNA recovery prior to further downstream analysis. Once best practices are established, a transition to full automatization of the sampling and preservation of filters/eDNA samples using remotely operated long-range mobile platforms and automated eDNA samplers could be adopted for medium- to large-scale biodiversity surveys (Preston et al., 2024)

### 4.2. Recommendations from this study

We encourage that the protocols regarding type of filters, sampling volume and sample conservation, are tested on a larger set of offshore samples prior to committing to best eDNA practices for these important steps of the eDNA workflow. Further, we recommend the following:

***Filters***

1. Sterivex filters are the recommended option for eDNA pretreatment. These filters reduce the risk of cross-contamination and are easier to handle than larger filters like Wattera. Furthermore, the biodiversity captured with Sterivex is comparable to that obtained with larger filters (Wattera) and disc filters (Durapore), making them a reliable choice for community analysis using metabarcoding.
2. For the targeted eDNA approach, for which the target is metazoan taxa, we suggest using Wattera filters. These filters offer greater surface area and higher filtration capacity, making them better suited for capturing the eDNA of rare or larger species as is demonstrated above.

***Preservatives***

1. Preservatives are not recommended for studies when aiming to assess whole community biodiversity using metabarcoding. Preservatives like SDS or BAC can alter community composition by introducing degradation of substrates. If preservatives are necessary, Longmire solution without SDS is a better option as it has less impact on eDNA-based outcomes. Optimal protocol and method for marine eDNA sample preservation is an area that deserves more research particularly in the offshore context where logistic and manpower capacity is limited.
2. For targeted detection and quantification of marine metazoans such as fish, preservative solutions added directly to seawater are more effective than freezing the filters.

