## Supplemental material and figures for "A practical evaluation of sampling filtration and preservation of environmental DNA samples for the water column monitoring"

**Running title: eDNA for use in offshore water column monitoring**

**Keywords**: offshore water column monitoring, environmental impact, eDNA sampling, eDNA conservation, digital PCR, eDNA metabarcoding

**Supplementary material**

**Supplementary TABLE 1**. Oligonucleotides used in this study.

| Type of analysis | Taxon of interest | Oligo name | Sequence (5' - 3') | Approximate product size (bp) | Reference |
| --- | --- | --- | --- | --- | --- |
| Metabarcoding | Prokaryotes | 341F  806R | CCTAYGGGRBGCASCAG  GGACTACNNGGGTATCTAAT | 460 | (Yu et al., 2005) |
| Metabarcoding | Eukaryotes | 528F  706R | GCGGTAATTCCAGCTCCAA  AATCCRAGAATTTCACCTCT | 220 | (Cheung et al., 2010) |
| dPCR | Dinophyceae | Dino681F  Dino723 (PROBE)  Dino809R | CTGTCAGAGGTGAAATTCTTGGA  TGCGAAAGCATTTGCCAAGGATGT  CTCTAGTCGGCATGGTTTATGG | 150 | (Krolicka et al., 2020) |
| dPCR | Teleostei | tele02_fw  tele02_rv | AAACTCGTGCCAGCCACC  GGGTATCTAATCCCAGTTTG | 167 | (Taberlet et al., 2018) |

**Supplementary FIGURE 1.** A) Total DNA (ng) yield recovered from Durapore (n=6), Sterivex (n=3) and Wattera (n=6) filters following filtration of the samples and DNA yield after normalization to 1L (ng/L). B) DNA yield normalized to 1L (ng/L) recovered from the different conservation solutions used for the water. Longi NoSDS- water samples preserved using Longmire solution without SDS, Longi- water samples preserved in original Longmire solution, BAC – samples preserved using BAC and filters Frozen –filter samples collected on Durapore filters and stored at -20°C on the ship then -80°C in the laboratory (see table1 in manuscript)

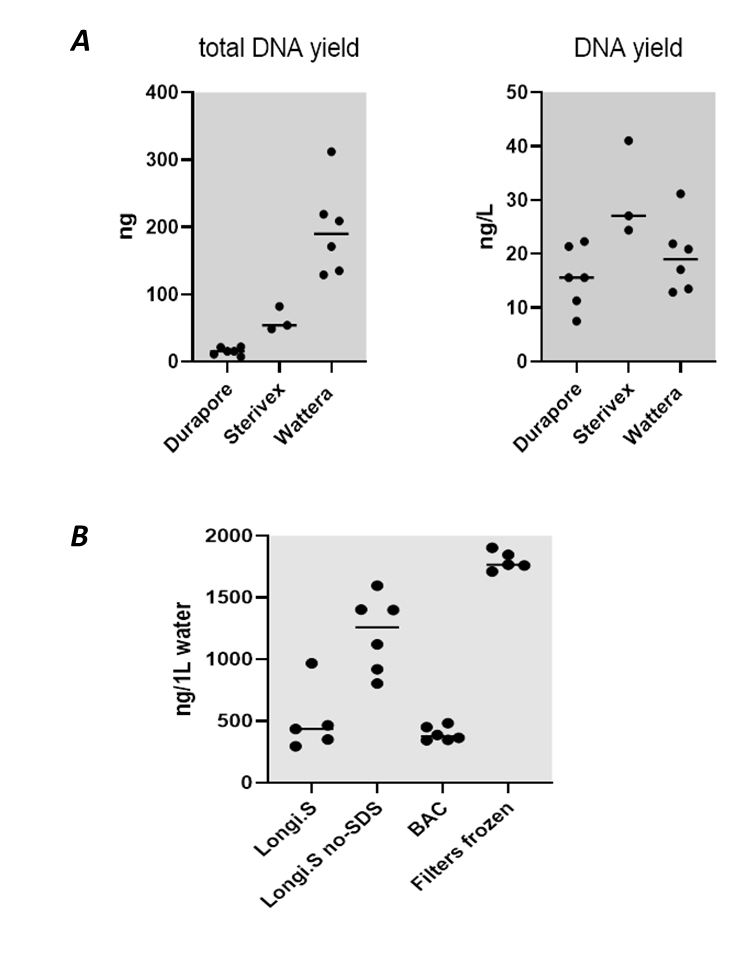

**Supplementary FIGURE 2.** Heatmap displaying different abundance and taxa clustering of the top 35 taxa (genus level) within *Prokaryotes* (A) and *Eukaryotes* (B) resulting from the use of the different filter types. Station3Ste, samples filtered with Sterivex; Station3Dur, samples filtered with Durapore; Station3Wat, samples filtered with Waterra

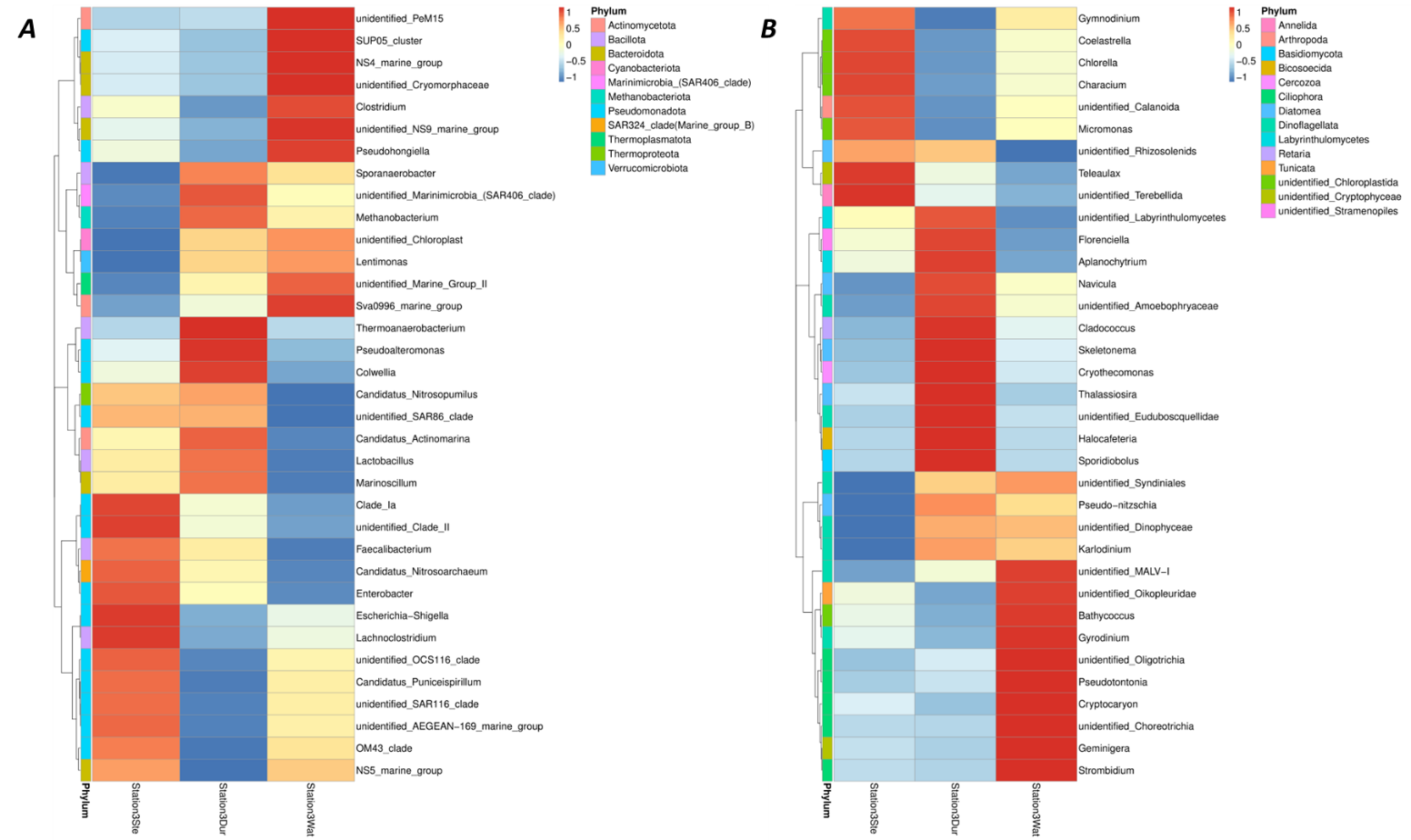

**Supplementary FIGURE 3 -** Histogram of the LDA scores for Prokaryotes (A) and Eukaryotes (B) following different filter types. Each bar represents species(biomarker) whose abundance shows significant differences among groups. The selecting criteria for significance is that LDA scores are larger than the set threshold (4 set by default). The length of each bar, namely, the LDA score, represents the effect size (the extent to which a biomarker can explain the differentiating phenotypes among groups). Station3Ste, samples filtered with Sterivex; Station3Dur, samples filtered with Durapore; Station3Wat, samples filtered with Waterra

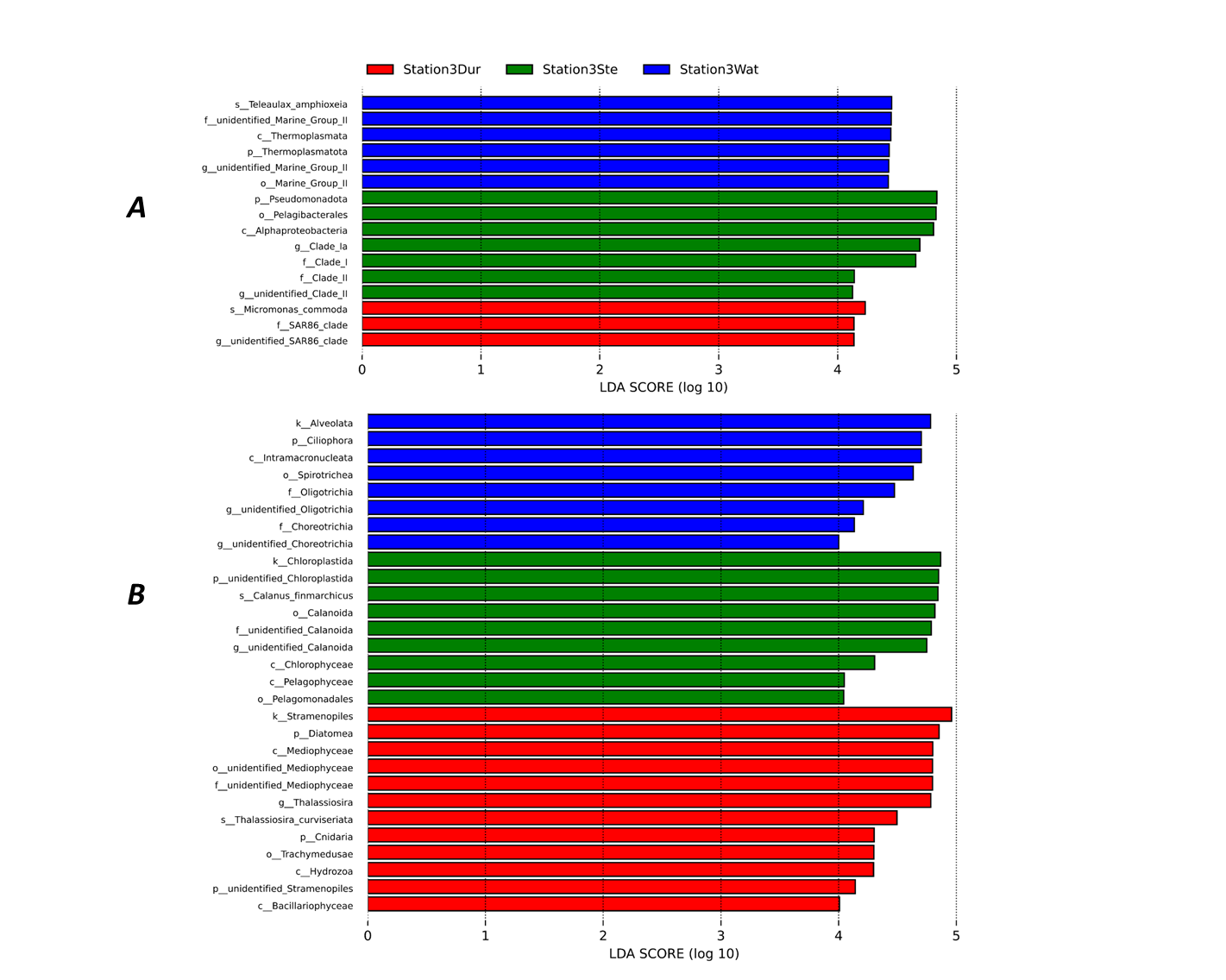

**Supplementary FIGURE 4.** Heatmap displaying different abundance and taxa clustering of the top 35 taxa (genus level) within Prokaryotes (A) and Eukaryotes (B) following different conservation options after 3 weeks. LNoSDS- water samples preserved using Longmire solution without SDS, LONGI- samples preserved with original Longmire solution, BAC – samples preserved using BAC and filtersFrozen – Durapore filters frozen at -20°C on the ship then -80°C in the laboratory

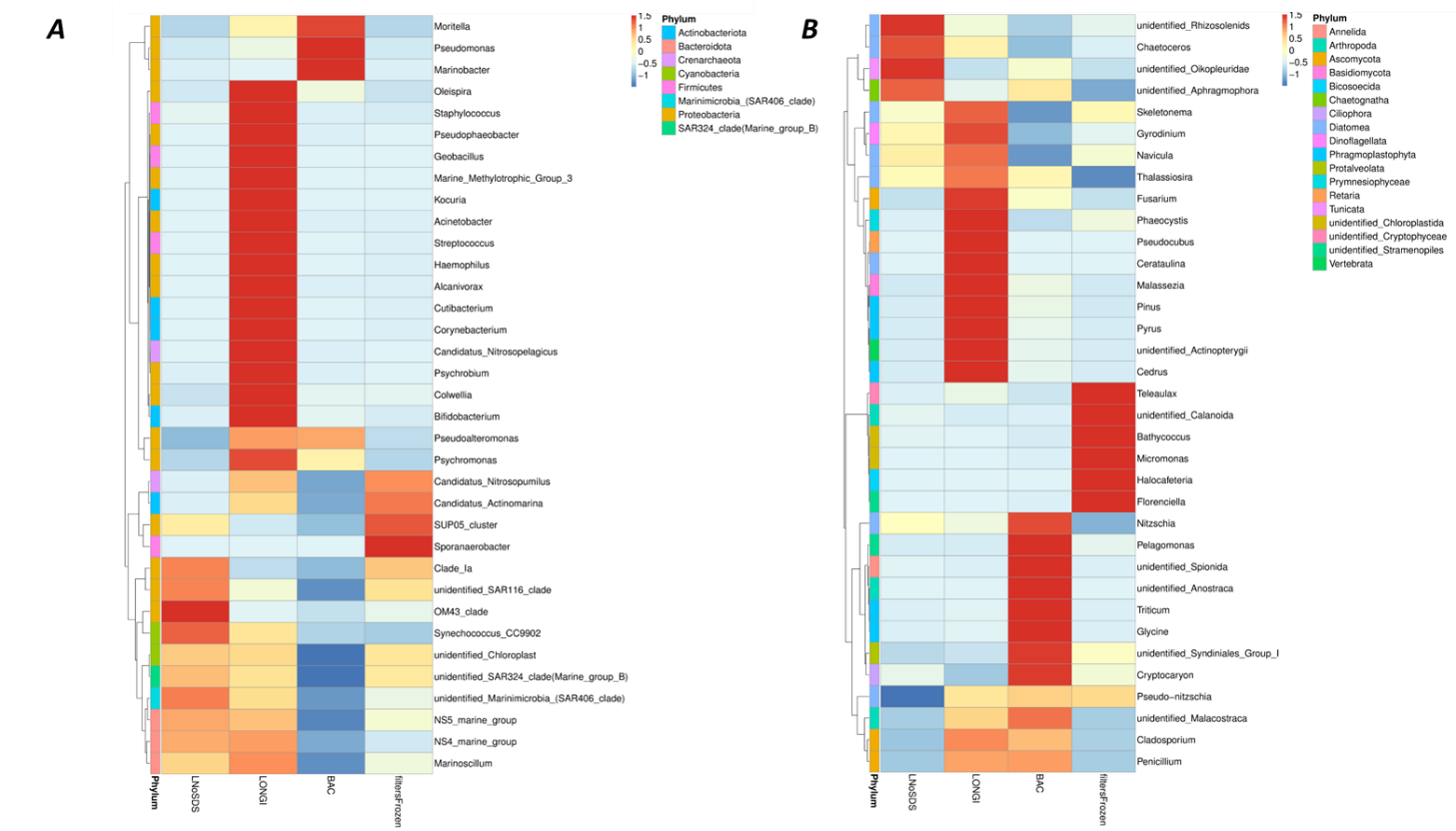
**Supplementary FIGURE 5 -** Histogram of the LDA scores for Prokaryotes (A) and Eukaryotes (B) following different conservation options after 3 weeks. Each bar represents species(biomarker) whose abundance shows significant differences among groups. The selecting criteria for significance is that LDA scores are larger than the set threshold (4 set by default). The length of each bar, namely, the LDA score, represents the effect size (the extent to which a biomarker can explain the differentiating phenotypes among groups). LNoSDS- water samples preserved using Longmire solution without SDS, LONGI- samples preserved with original Longmire solution, BAC – samples preserved using BAC and filtersFrozen – Durapore filters frozen at -20°C on the ship then -80°C in the laboratory

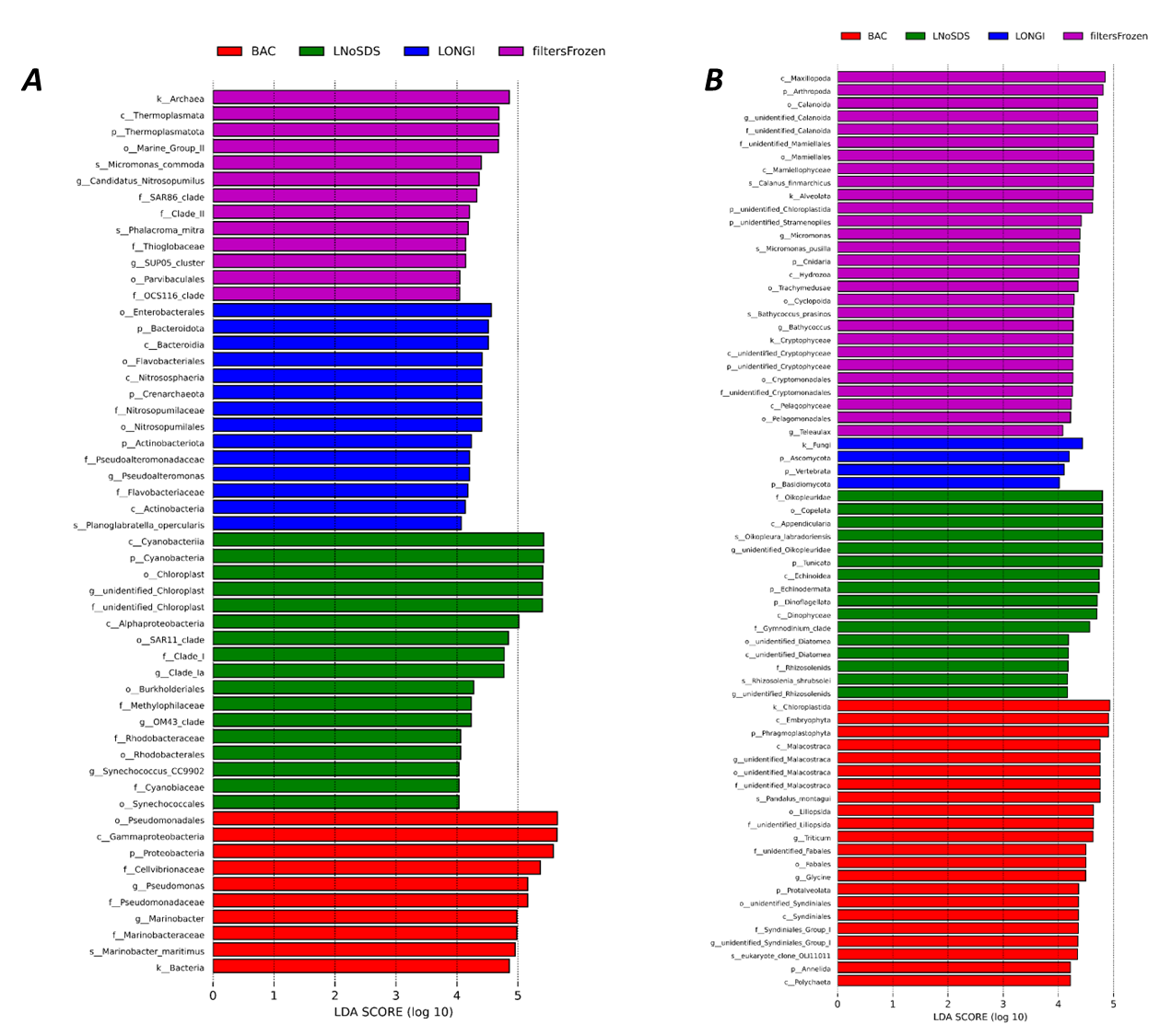
